# High-throughput chemical toxicity testing for beneficial insects: The value of coated vials

**DOI:** 10.1101/2025.02.02.636132

**Authors:** Joshua A. Thia, Julie Digard, Ashritha P.S. Dorai, Courtney Brown, Ary A. Hoffmann

## Abstract

Non-target effects of insecticides used in agriculture can impact the ecosystem services provided by beneficial insects. Understanding the broader effects of chemical usage requires multi-species investigations on the impact of different insecticide active ingredients. In this work, we tested the utility of coated vials as a quick, cheap, and efficient dried residue chemical toxicity assay. This method was compared against dishes sprayed with a Potter tower, which is an industry standard instrument used to apply precise insecticide doses. Our study focused on 2 natural enemies of aphids, larval and adult *Hippodamia variegata* ladybird beetle, and the parasitoid wasp, *Diaeretiella rapae*. These natural enemies were exposed to multiple doses of 3 insecticides that are registered for aphid control in Australian agriculture: alpha-cypermethrin (pyrethroid), dimethoate (organophosphate), and sulfoxaflor (sulfoximine). Modelled dose-response relationships showed evidence of general differences in the intercepts between methods, but not in slope (rate of mortality with concentration). However, these subtle statistically significant effects did not translate into marked differences in LC50-values, which were comparable across assay methods for all natural enemies and insecticides tested. Coated vials therefore produced estimates of chemical toxicity that were consistent with those derived from Potter tower sprays. This underscores the validity of coated vials as a method for performing high-throughput chemical toxicity experiments, as required when examining different populations of multiple taxa across a range of insecticide active ingredients.

## INTRODUCTION

Beneficial insects provide important ecosystem services in agricultural systems, including pest control, pollination and soil substrate development (Losey and Vaughan, 2006; Power, 2010). These services are, however, impacted by insecticides used for chemical pest control, which can have unintended acute (mortality) and chronic (sublethal) toxic effects on beneficial insects (Cole et al., 2010; Fernandes et al., 2016; McDougall et al., 2024; Overton et al., 2023; Skouras et al., 2019). Understanding the magnitude of chemical toxicity is crucial in developing guidelines for integrated pest management, supporting sustainable and more environmentally friendly agriculture (Schmidt-Jeffris, 2023; Serrão et al., 2022). By understanding which insecticides are ‘soft’ on beneficial insects, and ‘hard’ on pests, growers and agronomists can make more informed decisions about how on-farm practices might impact the ecosystem services they receive. Developing clear insight into how different insecticidal modes of action impact beneficial insects is becoming increasingly important as societal expectations shift to lowering chemical inputs into agriculture (Blanco-Moreno et al., 2024; Möhring et al., 2020; Sharma et al., 2019).

Drawing broad generalisations about the toxicity of different insecticidal modes of action is challenging. Multi-species studies of insecticide toxicity show that there can be considerable taxonomic variability in responses (Fernandes et al., 2016; McDougall et al., 2024; Overton et al., 2023). Such variability means that multiple species should be tested when developing generalizations. Unfortunately, chemical toxicity assays are typically labour intensive and can rapidly scale to an enormous size when many species, populations, insecticides, and concentrations are used, or when different responses are measured. Even the most expansive studies must trade off in some of these dimensions. For example, the focus may be restricted to many species but tested for a single active ingredient (Blanco-Moreno et al., 2024), many species and active ingredients but over a limited concentration range (Bernard et al., 2010; Overton et al., 2023), or many active ingredients and different responses but for a limited number of species (Gandara et al., 2024; Knapp et al., 2024; Lucas et al., 2004). Methods that allow researchers to generate meaningful toxicity data easily, quickly, and cheaply will help to broaden and deepen our understanding of insecticide impacts on beneficial insects.

All laboratory-based assays are artificial, in the sense that they do not fully capture the complexity of field conditions, but they are important in generating baseline expectations of chemical toxicity (Bernard et al., 2010; Blanco-Moreno et al., 2024; Mata et al., 2024). Many assay methods have been developed, and can vary largely among species and insecticides tested, as well as the types of responses measured (Bernard et al., 2010; Cole et al., 2010; Fernandes et al., 2016; Skouras et al., 2019). For natural enemies, a list of recommended methods has been established by the IOBC/WPRS (International Organisation for Biological Control/West Palaearctic Regional Section) (e.g., Hassan et al., 1992, 1985; Sterk et al., 1999). To achieve precise insecticide dosages and exposure relative to field rates, many methods use a Potter tower spray system to apply insecticides to an exposure surface (Potter, 1952). While the Potter tower is an important tool, it does have some practical limitations. Potter towers are relatively expensive, which may pose a challenge if funding is restricted, potentially discouraging practitioners from generating toxicity data or leading to concerns about whether the data generated meets industry standards. The throughput of a Potter tower is also inherently constrained, as only a small number of replicates can be sprayed together, and there is a time cost associated with handling. Furthermore, when working with multiple insecticides and dosages, thorough cleaning between sprays is required, adding to the time and effort needed to complete experiments.

Whilst chemical toxicity studies of natural enemies tend to focus on controlled assays, methods applied to arthropod pests tend to be wider in scope to allow for high-throughput investigations, often targeting locally cheap materials (Berlinger et al., 1996; Kabir et al., 1993; Umina et al., 2024; Zhao and Grafius, 1993). This contrasting approach to bioassays seems to reflect the different aims of testing programs. For natural enemies, single strains or populations are typically used to broadly represent expected responses of a species at large, with an emphasis on precise and accurate measurement. In pests, many strains or populations are often tested to understand variation of insecticide tolerance or resistance, with throughput playing a major factor in assay methodology (for example: Broadbent and Pree, 1997; Kwon et al., 2015; Thia et al., 2022; Zimmer and Nauen, 2011).

Coated vial assays provide a simple, quick, and affordable way to generate chemical toxicity data and have been applied to many arthropod pests (for example: Broadbent and Pree, 1997; Kwon et al., 2015; Thia et al., 2022; Zimmer and Nauen, 2011). Increasingly, these assays have been adapted for use in natural enemies (for example: Amarasekare et al., 2016; Liu et al., 2016; Perrin et al., 2024; Wu et al., 2007). One particularly notable study used coated vials to generate dose-response data for natural enemy communities in 5 European countries, encompassing 91 populations, and a total of 54 different species (Blanco-Moreno et al., 2024). Although coated vial assays clearly provide a logistical advantage, there are crucial knowledge gaps in understanding how toxicity measurements obtained using this method compare to those using recommended Potter tower spray protocols. Application of insecticide in a coated vial assays is achieved by simply swirling then decanting solution to leave residues on the vial internal surface. The exact dosage of insecticide is potentially more uncertain relative to what can be achieved with a Potter tower. Do coated vials therefore differ significantly in how they expose natural enemies to toxicity? And would different conclusions be reached if studies used coated vial over Potter spray methods?

In this work, we compared coated vial and Potter tower spray methods for generating chemical toxicity data for natural enemies. We focused on 2 natural enemies of aphids: the larval and adult life stages of the ladybird beetle, *Hippodamia variegata*, and the adult life stage of the parasitoid wasp, *Diaeretiella rapae*. Both natural enemies were assayed against 3 active ingredients registered for aphid control in Australian agriculture: alpha-cypermethrin (a pyrethroid), dimethoate (an orgaphophosphate), and sulfoxaflor (a sulfoximine). Our study was stratified into 3 experiments. In one experiment, we trialed the coated vial method to generate putative dose-response ranges for these natural enemies. In a follow-up experiment, we used these dose-responses ranges to directly test differences in mortality between coated vials and Potter tower sprays. Finally, we performed a validation test of whether the type of vessel (dish versus vial) affected mortality incurred by different insecticide application methods. The results suggest that the coated vial method can produce useful estimates of chemical toxicity that are comparable and consistent with those derived from Potter tower sprays.

## MATERIALS AND METHODS

### Insect species

Our study used two species of natural enemies of aphid pests. The first was the ladybird beetle, *Hippodamia variegata* (Coccinellidae). This species has a Palaearctic origin but is now distributed worldwide (reviewed in Franzmann, 2002). It has four life stages, and both larvae and adults are predators of aphids, but they also consume other prey species, such as mites, thrips, whiteflies and mealybugs (reviewed in Sarkar et al., 2023). Our second natural enemy was the parasitoid wasp, *Diaeretiella rapae* (Braconidae). This species has a putative west-Palaearctic origin but is also distributed worldwide (Carver and Starý, 1974). It parasitizes a range of aphid species that commonly infest brassica and grain crops (Ward et al., 2021). We used newly emerged *H. variegata* larvae (≤24 h post-emergence), adult *H. variegata*, and newly emerged adult *D. rapaae* (≤24 h post-emergence). This design captured two very distinct insect orders (Coleoptera versus Hymenoptera) and different life stages within *H. variegata*.

All natural enemies were sourced from the Biological Services (Loxton, South Australia 5333), a commercial biological control company. To prepare *H. variegata* larvae for our experiments, delivered eggs cards were broken into small pieces to isolate clusters of eggs. Each piece of egg card was placed into a 60 mm diameter Petri dish, sealed with parafilm, and incubated at 20°C until emergence. Adult *H. variegata* were delivered as post-emerged adults in small cardboard containers. Containers were held at 10°C for 2 days prior to assays. To prepare *D. rapae*, delivered mummies (dead aphids harbouring pupating wasps) were divided among small plastic cups and placed inside a larger plastic container, which was incubated at 20°C until emergence.

We also maintained a colony of the aphid species, *Myzus persicae*, as food for *H. variegata* in our assays. Our colony is a clonal isofemale line derived from a field strain in Victoria, Australia. This strain is raised on bok choy and maintained at a constant 20°C with a light:dark photoperiod of 16:8 hours.

### Insecticides

We measured toxicity to three insecticides active ingredients that are registered for aphid control in the Australian grain and horticulture industry: (1) Alpha-cypermethrin (Astound^®^ at 100 g/L, Nurfarm), a pyrethroid (IRAC group 3A); (2) Dimethoate (Dimethoate 400^®^ at 400 g/L, ADAMA), an organophosphate (IRAC group 1B); and (3) Sulfoxaflor (Transform^®^ 240 g/L, Corteva Agriscience), a sulfoximine (IRAC group 4C). For context, common recommended field rates (100L/ha) for aphid control in Australian are 125 mg a.i./L alpha-cypermethrin (grains), 300 mg a.i./L dimethoate (grains, legumes, and vegetables), and 240 mg a.i./L sulfoxaflor (legumes, pasture, and cotton). For all assays, insecticides were diluted in distilled water with 0.01% Tween (polysorbate) as a surfactant.

### Bioassay methods

Our study used three different methods to expose natural enemies to different insecticides: coated vials, coated Petri dishes, and Potter tower sprayed dishes. These are all ‘dried residue’ assays: liquid insecticide is applied to an exposure surface and allowed to dry, then insects are put in contact with these treated surfaces. The response recorded is the acute mortality observed in a set timeframe for the assay. Our assays used 30 mL plastic vials produced by ToolCraft Pty. Ltd. (Australia), 30 mm plastic Petri dishes produced by Greiner Bio-Oine (Austria), and 60 mm plastic Petri dishes produced by Corning Inc. (USA). We used plastic vessels as our contact surface because they were cheap and disposable, circumventing the need for decontamination between assays, as is needed for glass vessels.

Our coated vials method was similar to that described elsewhere for insect and mite pests (Broadbent and Pree, 1997; Kwon et al., 2015; Thia et al., 2022; Zimmer and Nauen, 2011). We coated the vials with insecticide by pouring solution into the vials, swirling the solution, then tipping it out. All natural enemies were exposed using the coated vial method. For our coated dish method, a 10 mL pipette was used to fill and remove insecticide solution from 30 mm plastic Petri dishes. Only larval *H. variegata* were exposed using the coated dish method. Our Potter tower spray method was similar to that described elsewhere for natural enemies (Knapp et al., 2024; Mata et al., 2024; McDougall et al., 2024; Overton et al., 2023), which followed IOBC/WPRS guidelines (Sterk et al., 1999). In our assay, Petri dishes were sprayed with 1 mL of insecticide solution at the equivalent rate of 100 L/ha using a Potter tower (Burkard Manufacturing Co. Ltd., Rickmansworth, UK). This is the typical rate used for many agriculturally important insecticides.

Calibration was performed with distilled water. All natural enemies were exposed using the Potter tower method. For larval *H. variegata* and adult *D. rapae*, we used 30 mm dishes, and for adult *H. variegata*, we used 60 mm dishes. Potter tower sprays were performed in replicate pairs of dishes for larval *H. variegata*and *D. rapae* assays, but dishes were sprayed individually for the adult *H. variegate* assays. Controls comprised water with 0.01% Tween. For all methods, insecticides were air-dried for several hours before storage at 1−3°C, with their lids on, for up to 4 days. Previous studies have stored coated vials or sprayed dishes for weeks at time without detectable effects on toxicity (Kabir and Chapman, 1997; Zimmer and Nauen, 2011), so we did not expect storage of treated vessels to be a caveat of our experiments.

Assays were always performed at least one day after preparing vials or dishes for logistical reasons. For adult *H. variegata*, we always performed assays 2 days after preparing plastics. For larval *H. variegata* and adult *D. rapae*, emergence was staggered over several days. This meant we also staggered the assay to use freshly emerged (≤24 hours) insects over consecutive days. For each day after plastic preparation, we checked the number of emerged larval *H. variegata* and adult *D. rapae*. If we could fill at least 6 replicates for larval *H. variegata*, or 3 replicates for adult *H. variegata* and *D. rapae*, *for* each insecticide, at each concentration (including controls), we then proceeded with the assay for that day. If we could not fill three replicates, all emerged insects were discarded for that day. Using this staggered approach, the mean assay day (after preparation of plastics) for larval *H. variegata* was 1.7 with a range of 1 to 3 days, and the mean assay day for adult *D. rapae* was 2.1 with a range of 1 to 4 days.

Our approach to filling vials or dishes with insects were as follows. For larval *H. variegata*, a 2% agar cube (∼0.5 cm^3^) was placed onto the lid to provide moisture along with 4 aphid nymphs (∼3-day old *M. persicae*) as a food source. We added a single larval *H. variegata* to each vial or dish because they are prone to cannibalism. For adult *H. variegata*, we also placed a 2% agar cube (∼0.5 cm^3^) onto the lid but provided 4 adult aphids as a food source. We added 3 adult *H. variegata* to each vial or dish. For adult *D. rapae*, a 1% agar 20% honey cube (∼0.5 cm^3^) was placed onto the lid to provide both moisture and food. We added 2 or 3 adult *D. rapae* to each vial or dish, depending on the total number of emerged wasps that day. Lids were screwed into vials and dishes were sealed were parafilm. For each insecticide, a set of control replicates were filled first before filling the treated replicates. We filled treated replicates up the concentration gradient. Insects were incubated at 18°C for 72 hours. Individuals were scored dead if they did not respond when disturbed with a fine horsehair paint brush, or if they could not right themselves when flipped on their back.

Note that we performed our assays over multiple runs for different experiments. Each assay comprised a unique batch of insects. We use letter codes A, B, C, D, E, F and G, to represent unique run IDs. Note that these run IDs are not transferable across the natural enemies studied. For example, Run A for larval *H. variegata* was performed on a different day to Run A for adult *H. variegata* and *D. rapae*. Summaries of sample sizes for each natural enemy, experiment, and run can be found in Supplementary Appendix 1.

### Experiment 1: Coated vial trials

Our first experiment used the coated vial method in trial runs to derive dose-response ranges for each natural enemy. These trials were used as proof-of-concept that coated vials could quickly and efficiently generate toxicity data for natural enemies. This data then informed our comparisons of coated vials with Potter tower sprays (see Methods ‘Experiment 2’ section below).

For larval *H. variegata*, we performed 3 independent runs, with 6 concentrations for Run A, 4 concentrations in Run B, and 5 concentrations for Run C, for each insecticide. Concentrations varied across runs as we tried to optimize the dose-response range for larval *H. variegata*. Depending on the emergence rate, each concentration was represented by 9, 10, or 15 replicate vials per run (Supplementary Appendix 1).

For adult *H. variegata*, we performed 2 independent runs, with 6 concentrations for Runs A and B, for each insecticide. The same concentrations were used in both runs for adult *H. variegata* because they produced good dose-responses. Each concentration was represented by 4 to 6 replicate vials per run (Supplementary Appendix 1).

For adult *D. rapae*, we performed 2 independent runs, with 6 concentrations for Runs A and B, for each insecticide. Concentrations varied between runs as we tried to optimize the dose-response range for adult *D. rapae*. Each concentration and control was represented by 6 replicate vials per run (Supplementary Appendix 1).

### Experiment 2: Comparison of coated vials and Potter tower sprays

Our second experiment tested the difference in dose-responses between coated vials and Potter tower sprayed dishes. Concentrations assayed for each natural enemy and insecticide were derived from the dose-response ranges in Experiment 1, specifically, the lethal concentrations expected to kill 20%, 50%, and 80% of individuals, the LC20-, LC50-, and LC80-values (see Methods ‘Statistical analyses’ section below). In principle, these 3 concentrations should be satisfactory to fit a dose-response relationship and estimate the LC50-value (Brown, 1978).

For larval *H. variegata* we performed 2 independent runs, Runs D and E, with 19 or 20 replicate vials or sprayed dishes for each concentration and control per run. For adult *H. variegata* we performed 2 independent runs, Runs C and D, with 5 replicates vials or sprayed dishes for each concentration and control per run. For adult *D. rapae*, we performed 2 independent runs, Runs C and D, with 6 replicate vials or sprayed dishes for each concentration and control per run. See Supplementary Appendix 1 for details.

### Experiment 3: Vessel effects

Our third experiment tested the interaction between vessel and insecticide application on mortality. The rationale for this experiment was that the vials and dishes used in this experiment represented quite different vessels in terms of their volume, surface area, and shape. Note that we were unable to establish the type of plastic used to make the vessels. Collectively, we wanted to test whether these combined effects could influence the mortality incurred by different insecticide applications (coating or Potter tower sprays). In our study, the coated dish method represents a hybrid method. It has the same volume and surface area of a Potter tower sprayed dish, but has insecticide applied using the pour-in-pour-out approach as a coated vial.

We performed Experiment 3 using just larval *H. variegata* and the LC50 concentrations for each insecticide. We performed 2 independent runs, Runs F and G, with 6 replicate vials or dishes for the control, and 12 or 18 replicates for treated vials or dishes, per run. See Supplementary Appendix 1 for details.

### Statistical analyses

All statistical analyses were performed in R. Observed mortality in treated replicates were corrected by the mean observed mortality in the controls using Abbott’s correction (Abbott, 1987):

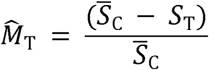

Here, 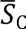 is the mean survival proportion observed in the control group, *S*_T_ is the survival proportion observed in a replicate, and 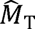 is the expected corrected mortality proportion in a replicate. Negative values were adjusted to zero. We performed Abbott’s correction in a nested fashion: replicates nested in run, runs nested in insecticide, insecticide nested in assay day.

Following Experiment 1, we performed initial analyses of dose-responses to generate LC20-, LC50-, and LC80-values for Experiment 2. We modelled mortality as a function of ‘Conc’, the fixed effect of the log10-transfromed insecticide concentration. We used the *glmmTMB* package (Brooks et al., 2017) to fit logistic regression models:

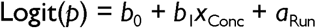

Here, the response is the logit-linked proportion of mortality, *b*_0_ is the intercept, *b*_1_ is the slope, *x*_Conc_ is the log-transformed insecticide concentration, and *a*_Run_ is the random intercept for run. We then used a modified version of the *dose.p* function from the *MASS* package to calculate the LC-values that informed concentrations tested in Experiment 2.

Before analysing dose-responses from Experiment 2, we tested whether replicates nested in a Potter tower spray could be considered independent or instead should be treated as paired replicates. We fit models for larval *H. variegata* and adult *D. rapae* separately and for each insecticide. In these models, we fit ‘Run’ as a categorical fixed effect, with levels ‘Run1’ and ‘Run2’. We also fitted the effect of ‘Conc’ as a categorical fixed effect, with levels ‘LC20’, ‘LC50’, and ‘LC80’. These were fitted as 2 logistic regressions using the *glmmTMB* package, taking the form:

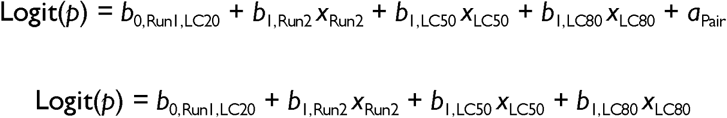

Here, the response is the logit-linked proportion of mortality. The *b*_0,Run1,LC20_ coefficient represents the intercepts for the reference level combination: ‘Run1’ and ‘LC20’. The *b*_1_ coefficients represent the deviations from the reference intercept based on whether a replicate was in ‘Run2’, and (or) exposed to the ‘LC50’ or ‘LC80’ levels of the insecticide. The *a*_Pair_ term is the random intercept for spray pair. The first model was the alternate model fitting the *a*_Pair_ term, contrasting to the second model which was the null model without the *a*_Pair_ term. We compared these models using the *anova* function, at a significance threshold of *P*<0.017 (0.05/3 insecticides). Because the *a*_Pair_ did not significantly explain more variance in the mortality responses (see Results), we proceeded with fitting our dose-response models without it. In other words, we could treat each sprayed dish as an independent replicate of concentration.

We then analysed data from Experiment 1 and Experiment 2 together. This allowed us to pool all our dose-response data together in a single model to compare methods and variance across different runs. However, for larval *H. variegata* and *D. rapae* in Experiment 2, we did not observe the expected mortality proportions for dimethoate assays – we therefore did not analyse these data (see Results). For all other natural enemy and insecticide combinations, we consider three different groups as levels of a fixed categorical ‘Method’ effect interacting with a fixed continuous ‘Conc’ effect. The levels of the ‘Method’ effect were: ‘Vial_trial’, the initial coated vial assays from Experiment 1, ‘Vial_test’, the coated vials assays tested against Potter tower sprays in Experiment 2, and ‘PT’, the Potter tower sprayed dish assays tested against coated vials in Experiment 2. This treatment of coated vials from Experiments 1 and 2 as separate groups also provided insight into the repeatability of toxicity assays using this method. We used *glmmTMB* to fit 3 logistic regression models, which took the form:

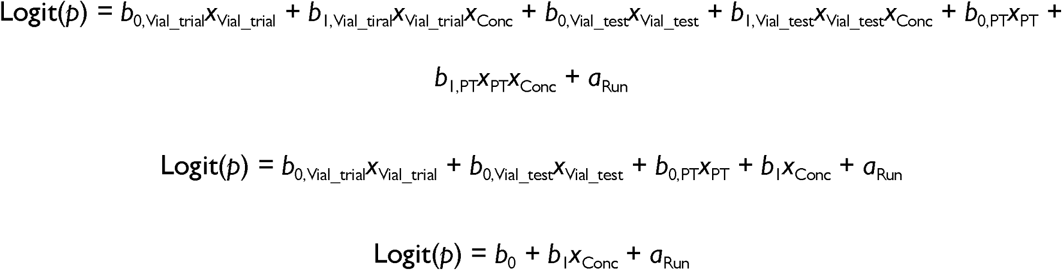

Here, the response is the logit-linked proportion of mortality. The *b*_0_ coefficients are the intercepts and *b*_1_ coefficients are the slopes, with group levels denoted by subscripts. The dummy variables *x*_Vial_trial_, *x*_Vial_test_, and *x*_PT_ are used to designate group levels, and *x*_Conc_ is the log10-transformed insecticide concentration. Random intercepts for each run are given by *a*_Run_. The first model is the full interaction model with separate intercepts and slopes for each group. The second model is an additive model with separate intercepts for each group but a global slope. The third model is a null model with a single global intercept and slope. These three models are hierarchical with decreasing complexity. We used sequential model comparisons with the *anova* function to determine the simplest model that could best describe the dose-responses. We first compared the interaction and additive model, and if these were non-significantly different, we compared the additive and null model. An analysis of deviance was performed on the best model using the *Anova* function from the *car* package.

Whilst our model comparisons were used to identify the simplest model, we were interested in how much estimated LC50-values varied between coated vials and Potter tower sprays, and between coated vials from the trials versus the test runs. We therefore estimated LC50-values using the full interaction models and performed post-hoc tests of the differences between pairs of methods:

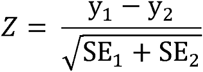

Here, the standardised difference, *Z*, is calculated as the difference between log10-transformed LC50-values, *y*, for groups 1 and 2, divided by the square-root sum of their standard errors. We obtained *P*-values for *Z*-scores using the *pnorm* function (assuming a mean of 0 and standard deviation of 1). We then designated statistical significance groupings using the *multcompView* package (Graves et al., 2019), correcting the statistical significance threshold of *P*<0.05 for multiple testing.

Finally, we analysed Experiment 3 to test the effects of vessel on pesticide application. In these analyses, we had a single fixed categorical factor, ‘Method’, with levels ‘Vial’ for coated vials, ‘Dish’ for coated dishes, and ‘PT’ for Potter tower sprayed dishes. We used *glmmTMB* to fit 2 logistic regression models:

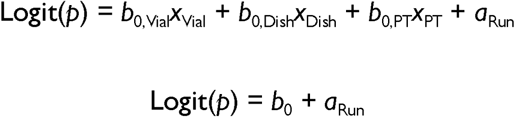

Here, the response is the logit-linked proportion of mortality. The *b*_0_ coefficients are the intercepts, with group levels denoted by subscripts, and *x*_Vial_, *x*_Dish_, and *x*_PT_ are their respective dummy variables. Random intercepts for each run are given by *a*_Run_. The first model was the alternate model fitting separate intercepts for each group, contrasting to the second model which was the null model fitting a single global intercept. We compared these models using the *anova* function, at a significance threshold of *P*<0.017 (0.05/3 insecticides). Similar to the dose-responses above, whilst our model comparisons provided a test of the simplest model, we wanted to compare differences between methods. We therefore used the *emmeans* package (Lenth, 2025) to perform post-hoc pairwise comparisons of mean mortality proportions estimated from the alternate models. We then designated statistical significance groupings using the *multcompView* package (Graves et al., 2019), correcting the statistical significance threshold of *P*<0.05 for multiple testing.

## RESULTS

### Control mortality

Generally, control mortality was low across out experiments. For larval *H. variegata*, the overall mean for proportional control mortality in Experiment 1 and 2 combined was 0.082 (±0.007 SE), and in Experiment 3 it was 0.11 (±0.0002 SE). For adult *H. variegata*, the overall mean for proportional control mortality in Experiment 1 and 2 combined was 0.13 (±0.009 SE). For adult *D. rapae*, the overall mean for proportional control mortality in Experiment 1 and 2 combined was 0.04 (±0.006 SE).

Notwithstanding, we note that control mortality varied across different combinations of assay runs and insecticides. These are illustrated in Supplementary Figures S1 to S4 and reported in the Supplementary Appendix 2. Most of these were well below the standard threshold of 0.2 typically used in chemical bioassays of invertebrates. However, others were not. In Experiment 3, control mortality was consistently greater in the coated dishes relative to other methods (Supplementary Figure S4). There is no clear reason why this would be the case: the Petri dishes used were the same as those used for the Potter tower sprays, and the only difference was the way in which the control solution was applied. Aside from the coated dishes in Experiment 3, there were no other obvious patterns for control mortality.

This apparently stochastic distribution of control mortality is unlikely to have major bearing on our conclusions, given that levels of mortality were low overall. Moreover, we completed multiple runs of the assays so that any variation among test samples would be included as variation among runs.

### Potter tower spray pairs

Our preliminary analysis of the Potter tower mortality data indicated that dishes sprayed at the same time could be considered separate replicates. Note, no mortality was observed in the dimethoate Potter tower sprays for adult *D. rapae*, so these were excluded from analysis. The Comparisons of models with the random effect of spray pair did not produce a significantly better fit than models without the effect of spray pair (*P*>0.8) (see Supplementary Appendix 3). We therefore treated each dish as an independent replicate, irrespective of spray group, in all further analyses.

### Coated vials versus Potter tower sprays

Our results from trials in Experiment 1 indicated that coated vials could provide an effective method for generating dose-response ranges for all natural enemies examined (Figure 1 & Table 1). We then used these dose-response ranges in Experiment 1 to test for derived predicted LC20-, LC50- and LC80-values. These predicted lethal concentrations generally produced expected outcomes in subsequent assays in Experiment 2 where we compared coated vials against Potter tower sprayed dishes (Figure 1 & Table 1). The exceptions were the dimethoate assays for larval *H. variegata* and the adult *D. rapae*. For both these natural enemies, the dose-response exhibited a very sharp jump from zero (or near-zero) mortality to complete mortality (Figure 1). This makes it difficult to precisely formulate concentrations that capture intermediate mortalities in the dose-response range. Hence, we were unable to compare methods for responses of larval *H. variegata* and adult *D. rapae* to dimethoate.

**Figure 1.**
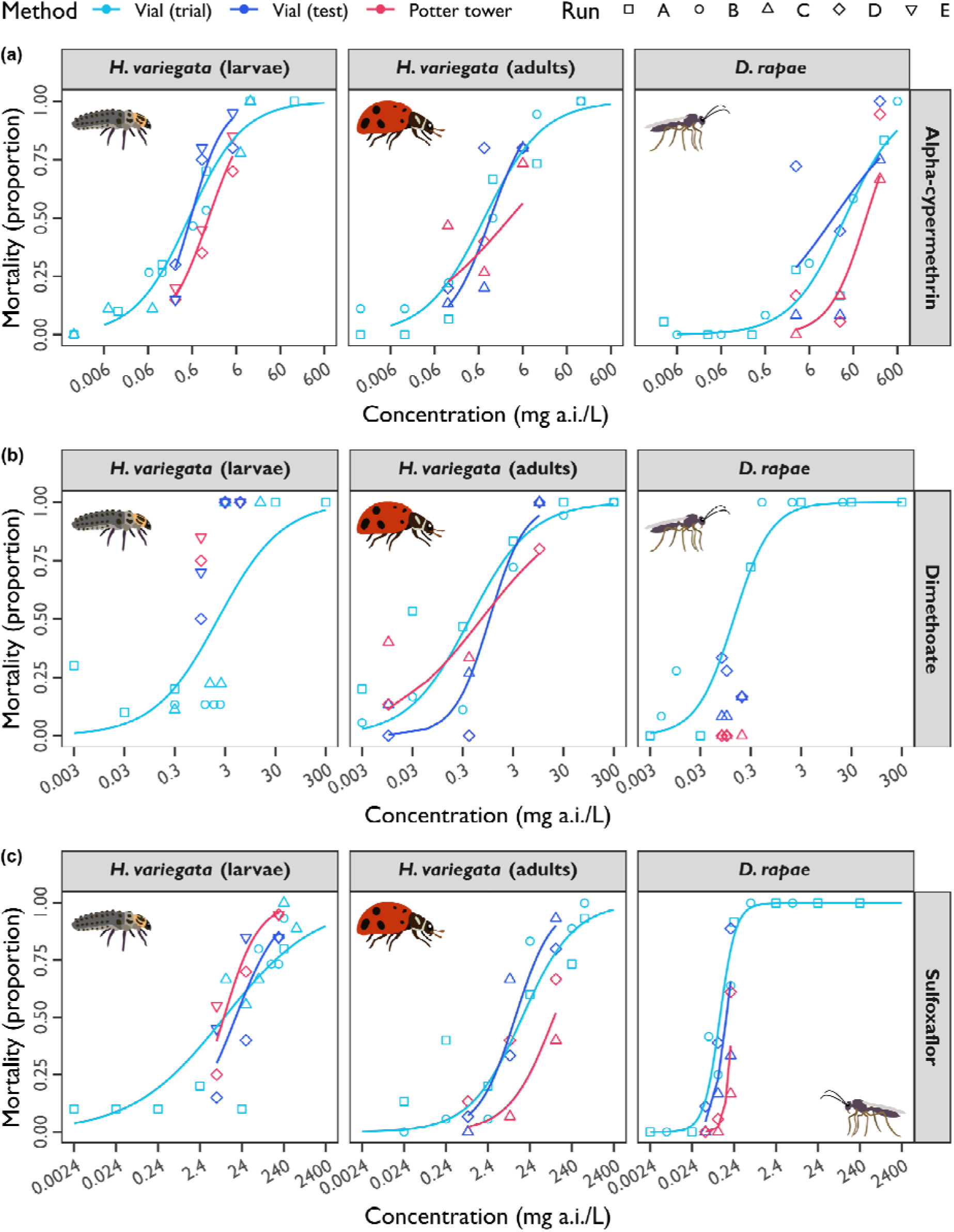
Dose-responses of natural enemies for different insecticides and exposure methods from fitted interaction models (Experiments 1 and 2). Concentrations of active ingredients are on the *x*-axis and mortality proportions are on the *y*-axis. Points represent observed mean mortality (Abbott’s corrected) with shapes indicating run and colour indicating method (see legend). Curves describe the predicted dose-response relationships with colour indicating method (see legend). Curves were not fitted if the range of mortality was not sufficient (that is, if proportions did not span 0.5). Panels contain data for each natural enemy (panel header) and different insecticides: (a) alpha-cypermethrin; (b) dimethoate; and (c) sulfoxaflor.

**Table 1.**
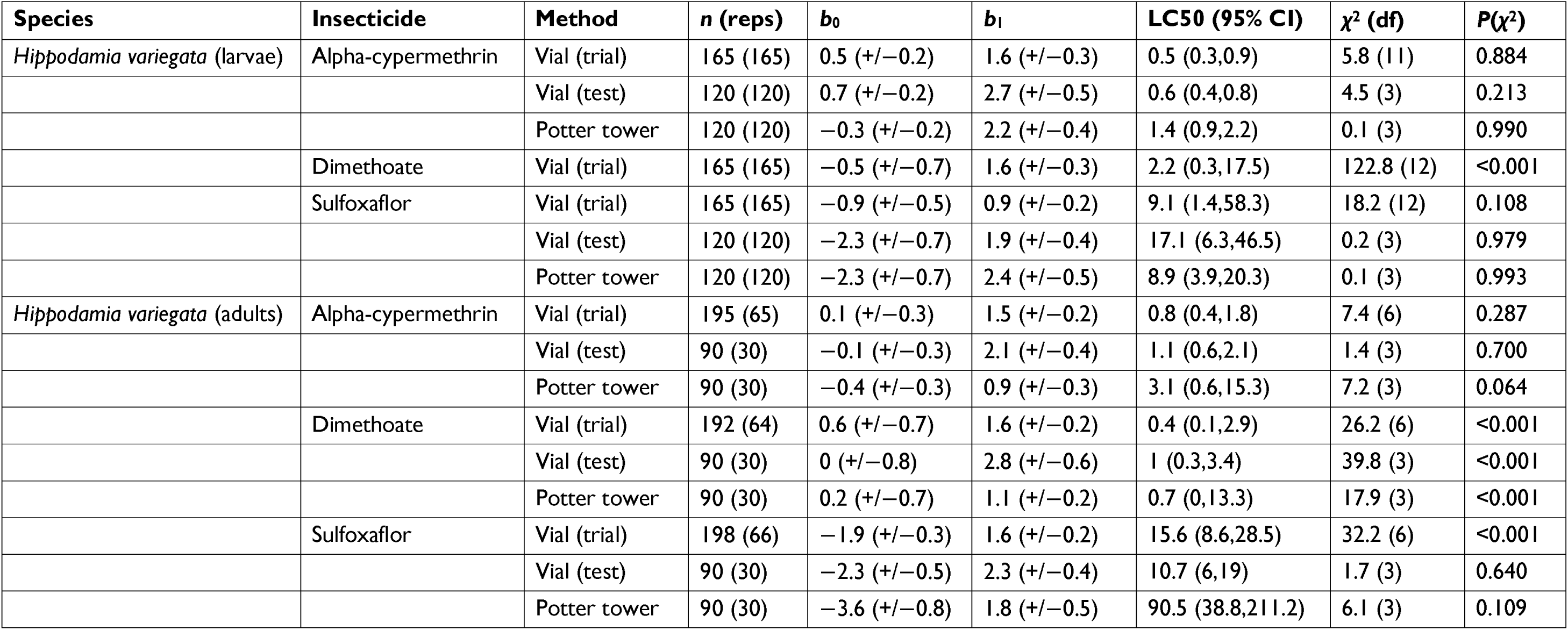
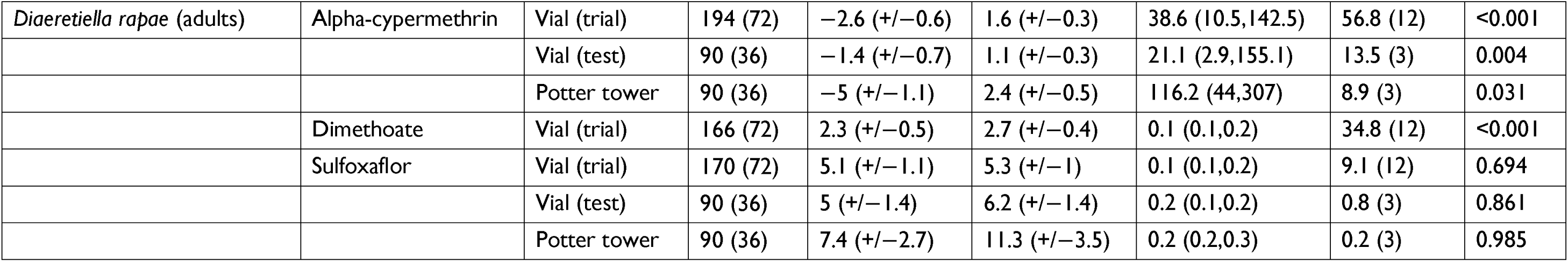
Dose-response statistics dervied from the interaction models for natural enemies for different insecticides and exposure methods from the fitted interaction model. Sample sizes are expressed as the total number of insects exposed (n) across independent replicates. Intercepts (b_0_) and slopes (b_1_) are reported on the log10-transformed mg a.i./L with standard errors. LC50-values and their 95% confidence intervals are reported in mg a.i./L. Goodness-of-fit for mean observed versus predicted mortality was assessed using χ^2^. Statistics were not calculated if the range of mortality was not sufficient, that is, if proportions did not span 0.5, as was the case for larval Hippodamia variegata and adult Diaeretiella rapae exposed to dimethoate (see also, Figure 1).

Our combined analysis of Experiments 1 and 2 tested the main effects of ‘Method’ (with levels ‘Vial_trial’, ‘Vial_test’, and ‘PT’) and ‘Conc’ (the insecticide concentration) and their interaction ‘Method:Conc’. Using hierarchical model comparisons (see Supplementary Appendix 4), the simplest models were: for larval *H. variegata*, the additive model for alpha-cypermethrin, and the interaction model for sulfoxaflor; for adult *H. varieta*, the null model for alpha-cypermethrin, the interaction model for dimethoate, and the additive model for sulfoxaflor; for adult *D. rapae*, the additive model for alpha-cypermethrin, and the additive model for sulfoxaflor.

The simplest model was therefore the null model in 1/7 comparisons, the additive model in 4/7 comparisons, and the interaction model in 2/7 comparisons. When the null model was the simplest, it suggests that all methods were comparable in their intercepts and slopes. When the additive model was the simplest, it suggests that methods varied in their intercepts, but not in their slopes. When the interaction model was the simplest, it suggests that methods varied in their intercept and (or) their slopes. In our case, however, the two significant interaction models only varied in their slopes, but not in their intercepts, for larval *H. variegata* exposed to sulfoxaflor and adult *H. variegata* exposed to dimethoate (Table 2).

**Table 2.**
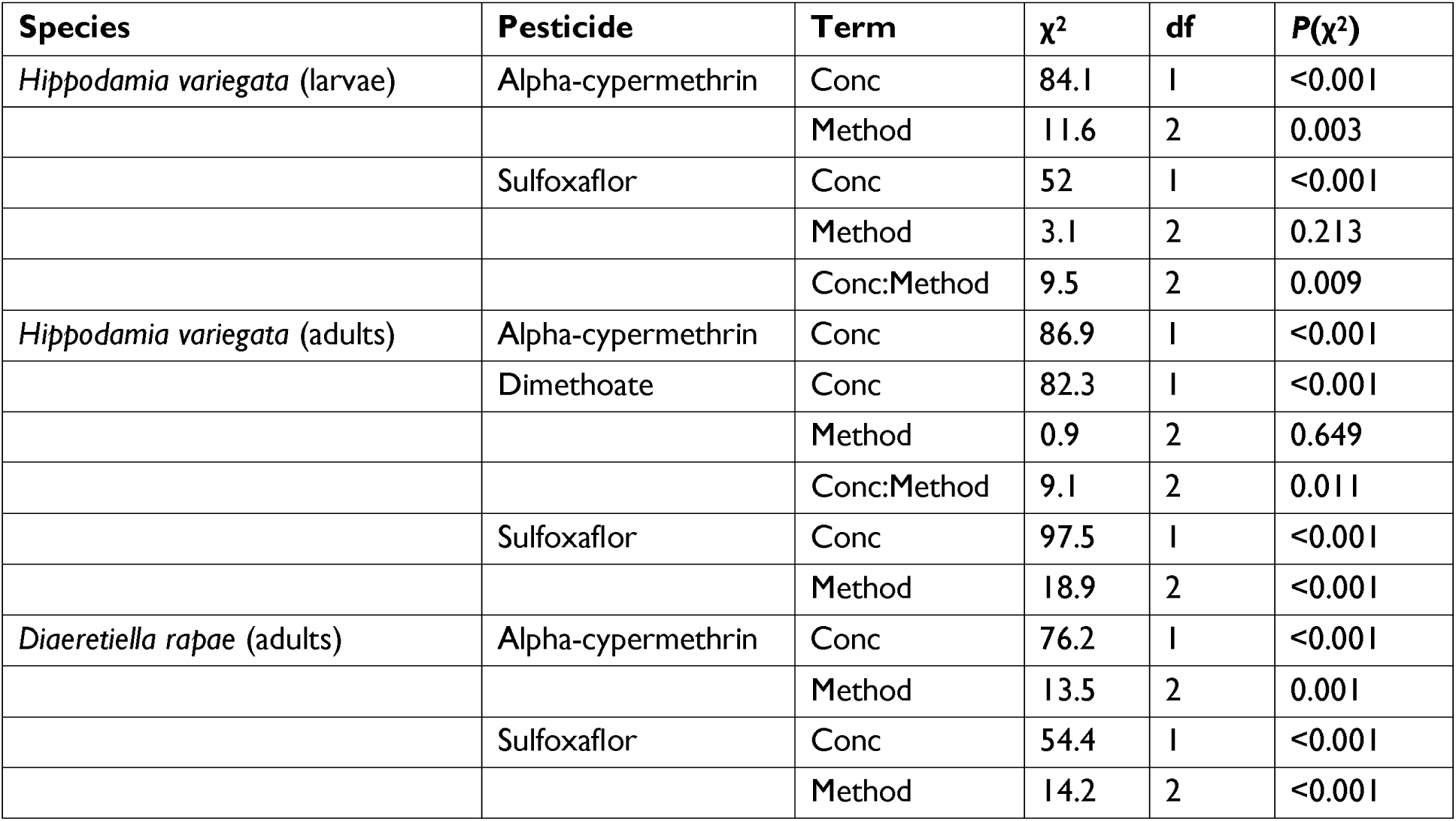
Analysis of deviance tables for the simplest dose-response models for natural enemies for different insecticides and exposure methods (Experiments 1 and 2). Statistics were not calculated if the range of mortality was not sufficient, that is, if proportions did not span 0.5, as was the case for larval *Hippodamia variegata* and adult *Diaeretiella rapae* exposed to dimethoate (see also, Figure 1).

Although our models suggested that significant variation in dose-responses could be attributed to method in some assays, this did not necessarily translate into marked differences in the estimated LC50-values. After correcting for multiple testing, almost all pairwise differences were statistically non-significant across insecticides and natural enemies (Figure 2 & Table 1). There was a tendency for Potter tower sprays to have slightly higher LC50-values, but the only statistically significant pairwise differences were for larval *H. variegata* exposed to alpha-cypermethrin and for adult *H. variegata* exposed to sulfoxaflor (in both cases, Potter tower sprays > coated vials) (Figure 2). By analysing coated vials from Experiments 1 and 2 as separate groups in our models, we were also able to demonstrate the consistency of estimates produced by this method. No pairwise differences in LC50-values were statistically significant between trial versus test coated vials (Figure 2).

**Figure 2.**
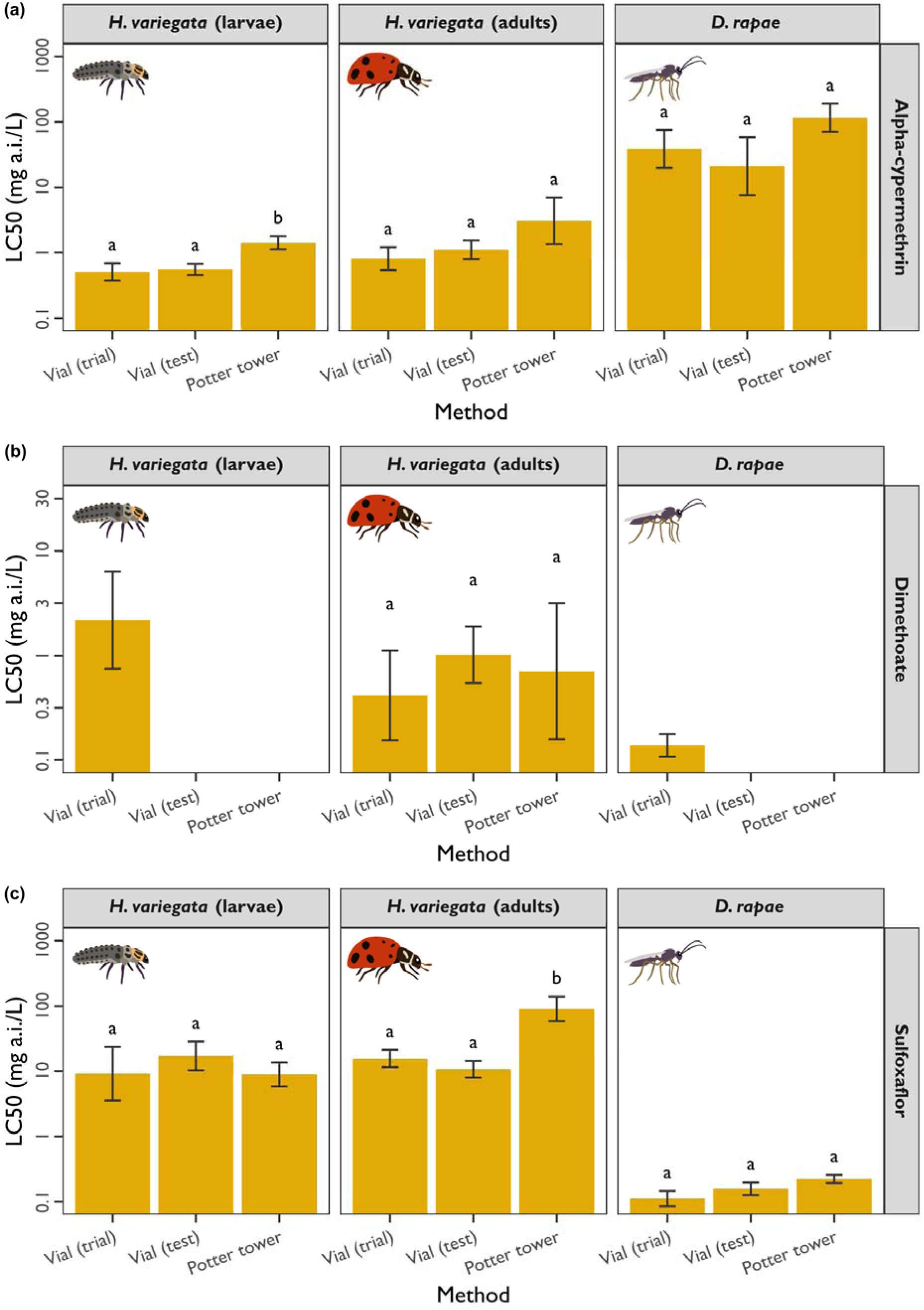
Post-hoc comparisons of LC50-values for natural enemies for different insecticides and exposure methods from fitted *interaction* models (Experiments 1 and 2). Methods are on the *x*-axis and estimated LC50-values (log10 scale) are on the *y*-axis. Gold bars represent the LC50 values estimates with 95% confidence intervals. Letters above points indicate significance groupings following corections for multiple testing. Note, significance groups were calculated across methods *within* natural enemies and should not be interpreted *among* natural enemies. Comparisons were not made if the mortality range was not sufficient, that is, if proportions did not span 0.5, as was the case for larval *Hippodamia variegata* and adult *Diaeretiella rapae* exposed to dimethoate (see also, Figure 1). Panels contain data for each natural enemy (panel header) and different insecticides: (a) alpha-cypermethrin; (b) dimethoate; and (c) sulfoxaflor.

### Vessel effects

Results from Experiment 3 indicated that that the different vessels used in this study are unlikely to contribute to differences in toxicity caused by insecticide application method. In this experiment using *H. variegata* larvae, we compared coated vials and Potter tower spray dishes to a hybrid coated dish method, at the predicted LC50-value for each insecticide. Hierarchical model comparisons (see Supplementary Appendix 5) indicated that the simplest model was the null model for the alpha-cypermethrin and dimethoate assays, and the alternate model was only favoured for the sulfoxaflor assay. In an analysis of deviance of the sulfoxaflor assay, the ‘Method’ term: χ^2^=9.1, df=3, *P*=0.028. However, after accounting for multiple testing, there were no significant pairwise differences in mean mortality for any of the insecticides (Figure 3).

**Figure 3.**
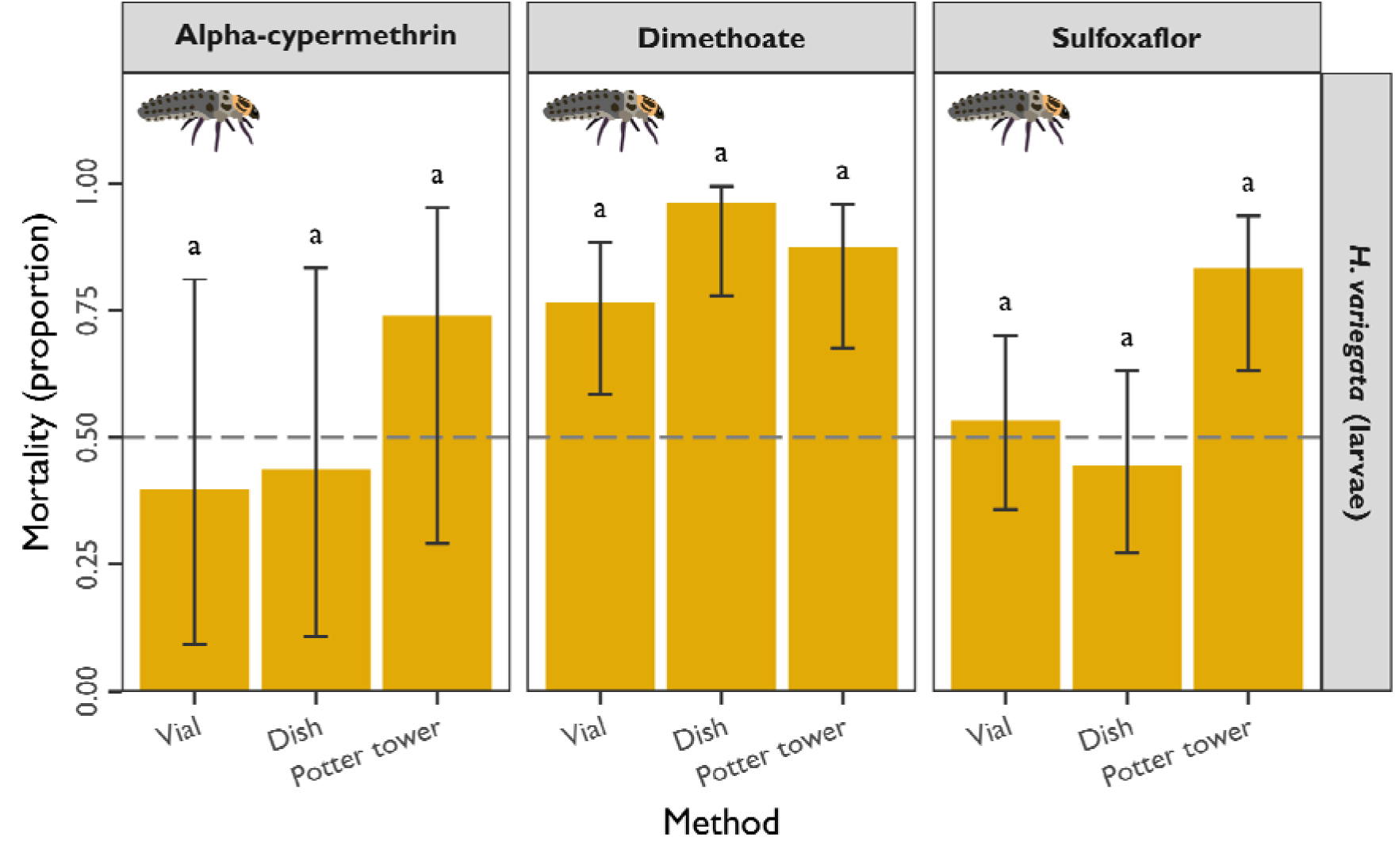
Estimated mortality of larval Hippodamia variegata exposed to insecticides using different applications and vessels from fitted alternate models (Experiment 3). Methods are on the x-axis and mortality proportions are on the y-axis. Gold bars represent the marginal means with 95% confidence intervals. Letters above points indicate significance groupings following corections for multiple testing. Grey dashed lines indicate mortality proportions of 0.5. Panels contain data for different insecticide active ingredients.

These assays for Experiment 3 also provided more evidence for the consistency of the coated vial method. For alpha-cypermethrin and sulfoxaflor, we were able to recover mortality that captured the predicted LC50-value (Figure 3). This was not the case for dimethoate (Figure 3), but as mentioned above, the sharp shift in mortality for this insecticide (Figure 2) may make it difficult to accurately formulate a concentration to achieve mortality at the target 0.5 proportion.

## DISCUSSION

Cost-effective and high-throughput chemical toxicity assays are useful for developing a broader understanding of how insecticides impact beneficial insects. Testing diverse insecticidal modes of action across multiple species is challenging but offers valuable insights into how beneficial insect communities might respond to commercially registered insecticides (Blanco-Moreno et al., 2024; Fernandes et al., 2016; Liu et al., 2016; Lucas et al., 2004). In this study, we demonstrated the value of coated vials for generating reliable toxicity data quickly, simply, and affordably for two natural enemies: *Hippodamia variegata* (ladybird beetle, larval and adult stages) and *Diaeretiella rapae* (parasitoid wasp, adult stage). Importantly, our results suggest that coated vials produced toxicity measurements consistent with those obtained using the Potter tower spray system, the industry standard for delivering precise insecticide doses to natural enemies. These findings are akin to Kabir et al. (1993) who undertook chemical toxicity assays in spider mites and showed that swirling insecticide in a Petri dish (their ‘PDR-R’ treatment) produced comparable mortality estimates as that with Potter tower sprays (their ‘PDR-PT’ treatment).

In our dose-response models, whilst there was a general trend for different intercepts, but not slopes, among methods, we did not observe major differences in predicted toxicity as inferred from LC50-values. Post-hoc tests of LC50-values showed that these estimates varied little among methods *within* natural enemies after accounting for multiple testing, irrespective of insecticide (Figure 2). *Among* natural enemies, we observed clear taxonomic variation in chemical toxicity that were consistent across methods: *H. variegata* was more tolerant to dimethoate and sulfoxaflor, and *D. rapae* was more tolerant to alpha-cypermethrin (Figure 2). By grouping the trial (Experiment 1) and test (Experiment 2) coated vial data separately, we also found consistency of this method across different runs. Repeated runs can help reduce the scale of single assays as well as capturing any stochastic variance in dose-response relationships. Consistency across methods was also reiterated in our follow-up experiment that confirmed that vessels did not interact with the insecticide application method (Experiment 3) (Figure 3).

A common feature of many chemical toxicity studies in natural enemies is the focus on single representative strains or populations (Mata et al., 2024; Overton et al., 2023). There are often understandable logistical reasons for this. Natural enemies can be much harder to rear than their pest hosts or prey, and field collected samples can be difficult to source in large numbers. Studies on natural enemies tend to prioritise suites of registered chemicals and (or) a range of different natural enemy taxa (Bernard et al., 2010). Our study also had this limitation, relying on a lab strain, whereas chemical tolerance can exist among field populations of natural enemies (reviewed in Hoy, 1990), and there may be opportunities to leverage natural variation to breed chemically tolerant natural enemies for commercial use (Balanza et al., 2021; Hoy, 1959; Liu et al., 2003). High-throughput methods, like coated vials, would be valuable for characterizing population variation, especially where the goal is to understand variation at expansive landscape scales (Blanco-Moreno et al., 2024).

While coated vial assays provide a useful screening tool where multiple tests are required, the Potter tower spray assays will no doubt remain the industry standard for topical applications of insecticide when delivery of a precise dose is necessary. Other methods for testing chemical toxicity will also remain important, with contrasting protocols and applications highlighting the complexity of interactions between species, insecticide active ingredients, and assay methods (Cole et al., 2010; Kabir et al., 1993; Lucas et al., 2004). But even when methods *within* species vary in their exact estimate of chemical toxicity, results may still be correlated *across* species, which is useful for comparative analysis (Wiles and Jepson, 1992). All laboratory-based assays are in essence artificial, and need to be validated in field trials (Fernandes et al., 2016; Fitzgerald, 2004; Schmidt-Jeffris, 2023; Umina et al., 2023), but they do provide crucial baseline data for comparing species and chemicals that inform practical recommendations (Bernard et al., 2010; Blanco-Moreno et al., 2024; Mata et al., 2024). The Potter tower remains an important instrument for studies aiming for field spray-like applications, direct contact of wet residues, or precise measured quantities of insecticide (Bernard et al., 2010; Desneux et al., 2006; Hassan et al., 1992, 1985).

In conclusion, we have shown the value of coated vial assays when compared to Potter tower assays by focusing on 2 important natural enemies and 3 core insecticides used to control aphids in Australian agriculture. There are growing issues with insecticide resistance in aphid pests in Australia and elsewhere (Morales-Hojas et al., 2020; Singh et al., 2021; Thia et al., 2024; Wang et al., 2018; Ward et al., 2024), highlighting the importance of biocontrol agents for controlling these pests and the need to expand chemical tolerance testing as new chemicals are deployed (Blanco-Moreno et al., 2024; Mata et al., 2024; McDougall et al., 2024). High-throughput assays therefore provide an important starting point for generating this vital data.

## ACKNOWLEDGEMENTS

We thank Biological Services for supplying the insects used in this study. We thank Evatt Chirgwin and Kathy Overton for providing training and advice in Potter tower usage. This work was part of an investment from the Grains Research and Development Corporation’s program, *Novel Pest Suppression in Grains and Vegetables: Preparing the grain and vegetable industries for a future without broad-spectrum insecticides* (UOM2404-006RTX), and Hort Innovation’s program, *Australian horticulture pest innovatio*(S*n*T23002). Funding for this work also came from the University of Melbourne.

## AUTHOR CONTRIBUTIONS

JAT designed the experiments, contributed to the bioassays, analysed the data, and wrote the original version of this manuscript. JD contributed to the bioassays and the original version of this manuscript. APSD contributed to the bioassays. CB contributed to the bioassays. AAH contributed to experimental design and obtained funding for this work. All authors contributed to the final version of this manuscript.

## DATA AVAILABILITY

All data and analyses required to replicate this study are available in a FigShare repository (Thia, 2025).

